# Diabetic liver-enriched secretory dipeptidyl peptidase 4 (DPP4) fuels gut inflammation via attenuation of autophagy

**DOI:** 10.1101/2024.08.06.606776

**Authors:** Mohammad Athar, Ratulananda Bhadury, Chayanika Gogoi, Pooja Mishra, Prity Kumari, Manisha Yadav, Jaswinder Singh Maras, Devram S. Ghorpade

**Affiliations:** Immuno-Inflammation Laboratory, National Institute of Immunology, New Delhi, India; Department of Molecular and Cellular Medicine, Institute of Liver and Biliary Sciences, New Delhi, India

**Keywords:** Diabetes, Hepatokine, DPP4, Autophagy, Colitis, Liver-gut cross talk

## Abstract

The recurrent pathological inflammation of the gut is a major concern in diabetic patients. With the failure of anti-inflammatory or diabetic drugs to limit relapse of colon inflammation demands the unearthing of mechanistic details underlying higher incidences of colitis in diabetic patients. Here we report the enrichment of DPP4 in the livers and blood samples of diabetic humans and mice models of diabesity that is in parallel to the development of colitis. Overexpression of DPP4 exacerbates or hepatic silencing of DPP4 impairs experimental colitis induced by DSS and STM. Mechanistically, we identified liver DPP4 attenuates gut-autophagic response to trigger enteric cell apoptosis, reduced mucin secretion, and compromised gut barrier leading to high infiltration of immune cells secreting inflammatory cytokines establishing pathological gut inflammation. Thus, liver-DPP4-mediated gut autophagy inhibition is a key pathway in diabesitic colitis.

## Introduction

Obesity-associated diabetes (diabesity) is a disease of disruptive energy homeostasis leading to the sub-optimum functioning of multi-organ system in the body(1). Low-grade, chronic inflammation during diabesity is thought to be at the roots of diabesity-associated pathological comorbidities including cardiovascular disease, fatty liver disease, kidney failure, inflammatory bowel disease (IBD), etc.(2,3). Diabesity and concomitant rise in a variety of comorbidities suggest the failure of effective communication among multi-organ system(4). Uncovering the cross-communication via plasma-soluble organokines is a less explored area of research. Recent studies have illustrated the inflammatory cross-talks among metabolic organs during diabesity. In diabesity, the liver is a major endocrine organ that adapts to disruptive energy homeostasis via secreting an array of hepatokines(5). For example, liver secretory Fetuin B (FETUB) triggers microglial inflammation causing diabetic retinopathy(6), dipeptidyl peptidase 4 (DPP4) instigates VAT inflammation(7), Fetuin A (FETUA) increases the secretion of monocyte inflammatory cytokines in adipose tissue(8). These studies summarize the core role of hepatokines in communicating with other organs and triggering pathological inflammation. However, no study has so far identified the role of hepatokines in a diabesity-associated increased incidence of gut inflammation.

IBD is sub-grouped into Crohn’s disease (scattered inflammation throughout gut) and colitis (inflamed colon)(9). Over the past decades, an increasing number of diabetic patients have been admitted to hospitals with severe abdominal pain with bloody stool(10,11). The colonoscopy observation indicated the excessive infiltration of neutrophils and lymphocytes with red patches throughout the colon confirming the development of colitis. Metformin therapy could provide a brief relief without complete remission of colitis(12). Current therapeutic regimes to treat diabesitic-related incidences of colitis fall short due to an incomplete understanding of the molecular events underlying pathological colon inflammation.

At a molecular level, colitis is characterized by reduced mucin protection, compromised gut barrier, apoptosis of epithelial cells, excessive infiltration of immune cells, and high secretion of inflammatory cytokines(13,14). One of the key aspects of developing colitis is a failure of gut homeostasis processes like autophagy(15). The process of maintaining gut health is governed by the autophagic response to invading pathogens, boosting an effective innate response, regulation of mucin secretion by enteric goblet cells, and promoting epithelial cell proliferation to counter pathological insults. A growing pieces of evidence suggest that defects in the autophagy process are closely linked to colitis(16,17). In corroboration, the induction of autophagy ameliorates experimental colitis, highlighting the crucial role played by autophagy in maintaining gut homeostasis. Despite a sufficient understanding of autophagy and its role in the pathophysiology of colitis, information on compromised autophagic response in diabesity is not well known. In particular cross-communication between liver-to-gu0t- and mediating forces is not well illustrated. In the present study we provide support to this idea and identify the role of diabesitic hepatokine in the attenuation of a gut-autophagy as a major cause of diabesity-related increase in incidences of colitis.

## Results

### Selective expression of DPP4 in metabolically disrupted human and mice livers positively correlates with gut inflammation

We aimed at metabolic syndromes that majorly affect the functioning of the liver like NASH, NAFLD and diabetes to identify unique secretory hepatokines **(Supplemental table 1)**(5). For this purpose, we looked into publicly available datasets and analysed them using GEO2R tool to identify differentially expressed hepatokines. We curated the expression profile of hepatokines in liver biopsies obtained from healthy donors (n=5) vs NASH patients (n=12), GSE24807; healthy donors (n=4) vs NASH patients (n=7), GSE17470; healthy donors (n=7) vs NASH patients (n=11), GSE63067; healthy donors (n=10) vs NAFLD patients (n=206), GSE135251; and NAFLD patients (n=51) vs NASH patients (n=47), GSE167523 (**Fig. 1A-E, Supplementary Fig. S1A-E’ and Supplementary Fig. S2A-T**). We also looked into secretory hepatokine profiles of diabetic patients in two separate human genomic datasets, GSE26168 (Healthy, n=8 vs DM, n=9) and GSE250283 (Healthy, n=15 vs DM, n=20). Although all the human datasets indicate the expression of a couple of hepatokines with little or no overlap with varying degrees of expression, to our surprise DPP4 is consistently upregulated in all of the human datasets obtained from metabolically deregulated livers of NAFLD and NASH patients (**Fig. 1A-F, Supplementary Fig. S1A-E’ and Supplementary Fig. S2A-T**). Circulatory DPP4 levels are elevated in obese-diabetic patients(18). Consequent data curation performed on blood samples obtained from DM patients suggested a marked augmentation of circulatory DPP4 levels (**Fig. 1A; 1G, H and Supplementary Fig. S2U-E’**). To corroborate with human datasets findings, we investigated the livers and plasma samples obtained from high-fat diet (HFD) fed and genetically obese-diabetic leptin receptor mutant (Lepr^db/db^ or db/db) mice models. Upon feeding of 60% kCal derived from a fat diet for 16 weeks, mice were obese with accumulation of visceral adipose tissue (VAT) mass. The cardinal feature of diabesity like hyperglycemia, hypercholesterolemia and glucose intolerance were documented in HFD-fed mice (**Supplementary Fig. S3A-F**). Similarly, leptin receptor mutant db/db mice on a regular chow diet were morbidly obese and diabetic (**Supplementary Fig. S3G-L**). Like human cohorts (above), we measured the liver expression of DPP4 levels in these mouse models of prediabetes (HFD-fed) and morbid diabesity (db/db). *Dpp4* transcript levels were markedly elevated in the liver of HFD-fed and db/db mice with concomitant accumulation of liver DPP4 (**Fig. 1I-N)**. Considering the high rate of IBD dominance in diabetic patients, we evaluated the gut inflammatory signatures in the mouse models of diabesity. Upon examination of colon tissue from chow-fed mice and HFD-fed prediabetic mice, we found a significant reduction in colon length, enhanced infiltration of immune cells into submuscularis and muscularis layer of colons with marked up-regulation of human colitis relevant inflammatory markers, *Mcp1, Il-6, Lcn2, Tff3, Camp, Il-1rn. Il-1ra, Pdgfb,* and *Crp* (**Fig. 1O-S**). Similarly, morbidly obese db/db mice presented with cardinal features of colitis like reduction in colon length, colonic over-expression of one of the inflammatory cytokines, MCP1 with enhanced expression of human colitis relevant inflammatory panel of genes (**Fig. 1T-W**). These data positively establish the co-occurrence of colitis in the settings of diabesity. To further establish the diabetic colitis phenotype and support the observation of human diabetic patients (with higher DPP4 levels) being vulnerable to developing colitis, we evaluated the correlation dynamics between liver DPP4 expression levels to that of colon inflammatory markers. We found a decreased colon length with incremental expression of liver DPP4. Conversely, Liver DPP4 levels are positively correlated with colitis markers, *Lcn2, Crp*, and *Tff3* in both HFD-fed and db/db mice models of diabesity (**Fig. 1X-A’ and Fig. 1B’-E’**). Therefore, these data support the idea of liver to gut cross communication possibly via secretory DPP4 and set the stage to validate the role of liver secretory DPP4 in triggering colon inflammation in the setting of diabesity.

**Figure 1.**
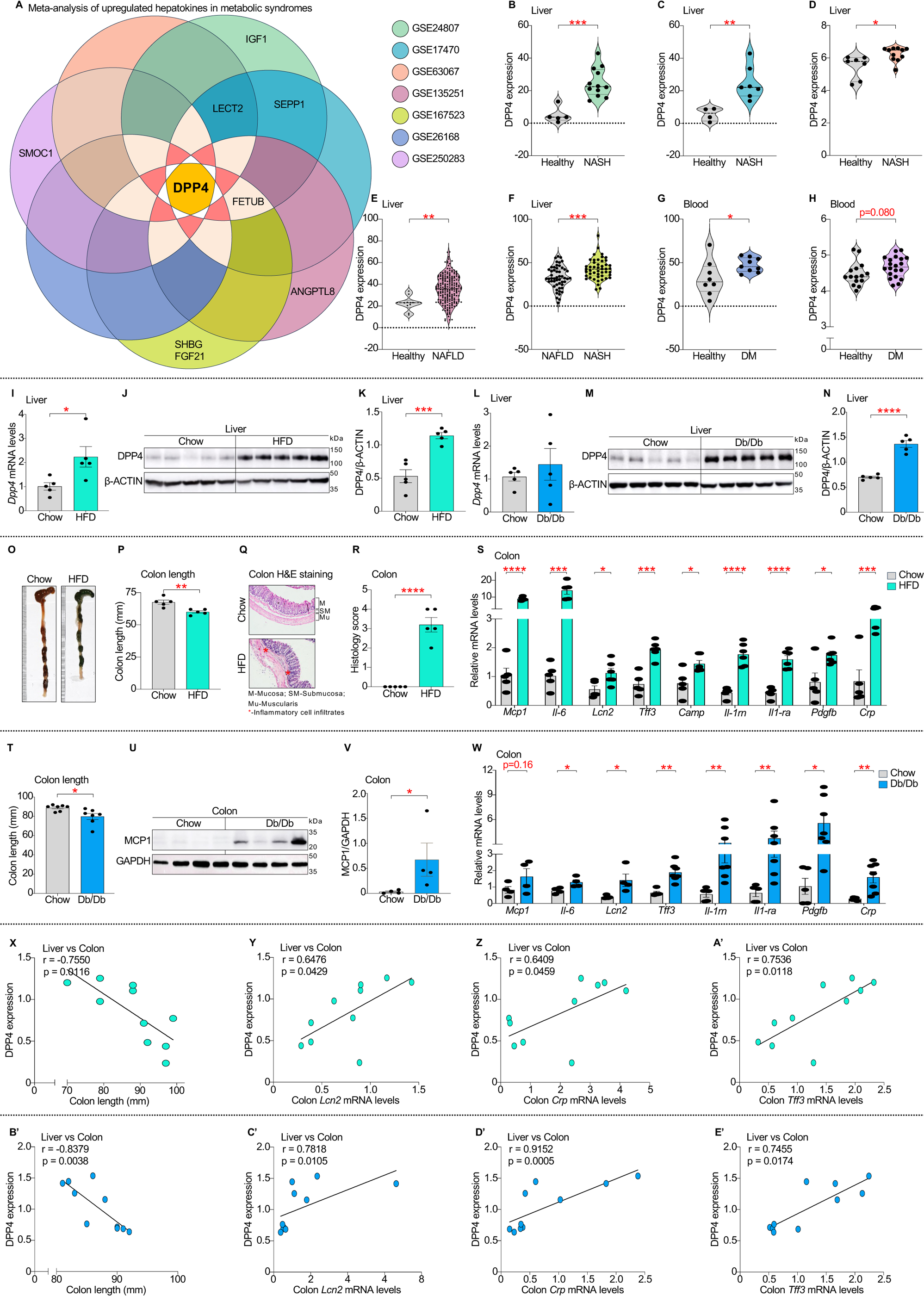
Marked upregulation of hepatic DPP4 in diabesitic humans and mice positively correlates with gut inflammation. **A**, Common and differentially upregulated hepatokines among 7 separate human genomic special event datasets, GSE24807, GSE17470, GSE63067, GSE135251, GSE167523, GSE26168 and GSE250283. **B-D**, DPP4 expression levels in the livers obtained from Nonalcoholic steatohepatitis (NASH) patients vs Healthy individuals (GSE24807; P < 0.001, GSE17470; P < 0.01, and GSE63067; P < 0.05). **E**, DPP4 expression levels in the livers obtained from Nonalcoholic fatty liver disease (NAFLD) patients vs Healthy individuals (GSE135251; P < 0.01). **F,** DPP4 expression levels in the livers obtained from NAFLD patients vs NASH patients (GSE167523; P < 0.001). **G & H**, Plasma DPP4 levels in diabetes mellitus (DM) patients compared to healthy individuals (GSE26168; P <0.05, and GSE250283; P =0.080). WT C57BL/6J mice were fed chow or HFD for 16 weeks and the following parameters were assayed. **I**, Liver *Dpp4* mRNA levels. **J & K**, Liver DPP4 protein expression, and quantification relative to GAPDH levels in chow vs HFD-fed mice. **L-N**, Liver *Dpp4* mRNA levels and liver DPP4 protein expression with quantification in WT and Db/Db mice. **O-S**, In Chow and HFD-fed mice colon pathology was assessed by colon images along with measurement of colon length (**O & P**), representative H&E-stained colon histopathology images along with histology score (**Q & R**) and colitis-specific expression of *Mcp1, Il-6, Lcn2, Tff3, Camp, Il-1rn, Il-1ra, Pdgfb,* and *Crp* (**S**). Wt and db/db mice colon inflammation is evaluated by **T**, colon length estimation, **U & V**, MCP1 levels in the colon with relative quantification by densitometry. **W**, *Mcp1, Il-6, Lcn2, Tff3, Il-1rn, Il-1ra, Pdgfb,* and *Crp* levels in the colon. **X-E’**, Correlative analysis between HFD-fed mice liver DPP4 levels and (**X**) colon length, (**Y**) colonic *Lcn2*, (**Z**) colonic *Crp*, and (**A’**) colonic *Tff3.* Analysis of correlation between db/db mice liver DPP4 levels and **(B’**) colon length, (**C’**) colonic *Lcn2*, (**D’**) colonic *Crp*, (**E’**) colonic *Tff3.* N = 4-8 mice per group; Data are mean ± SEM; ns, non-significant; *, < 0.05; **, < 0.01; ***, < 0.001; ****, < 0.0001 by unpaired t-test or by Mann Whitney t-test.

### Overexpression of DPP4 augments dextran sulphate sodium (DSS)-induced experimental colitis

To investigate the role of DPP4 in fueling colon inflammation, we first employed a DPP4-gain-of-function strategy. The circulatory DPP4 levels were upraised using adenoviruses carrying DPP4-overexpression construct. 10 days post injection of adeno-DPP4 viruses, we confirmed DPP4 over-expression in mice by measuring circulatory DPP4 activity (**Supplementary Fig. S4A**). The etiology of pathological colon inflammation during diabesity is complex with the gut being subjected to multifactorial insults that cumulates into pathological inflammation. To mimic these complexities and evaluate the contribution of DPP4 in these processes, we combined DPP4 overexpression with experimental colitogenic stimuli DSS. The effect of DSS insult on the inflammatory pathology of colon was evaluated by varying the doses of DSS and monitoring mice over a five-day time frame. We noted that a low dose of DSS (1%) induces suboptimum colitis as observed with kinetic analysis of percent decrease in body weight (BW), lowering of colon length, colon histopathology and expression of colonic inflammatory cytokines, *Mcp1* and *Il-6* (**Supplementary Fig. S4B-H**). Based on these data we selected a low-dose DSS (1%) model to investigate the regulatory role of hepatic DPP4 on colitis. We split the control cohort and Adeno-DPP4 cohort (DPP4 overexpression) into water and low-dose (1%) DSS cohorts. Five days post-DSS treatment, DPP4 overexpressing cohorts showed a marked reduction in colon length, higher colonic infiltration of immune cells, and elevated circulatory MCP1 levels compared to mice that were injected with control adenoviruses overexpressing GFP (**Fig. 2A-F**). We observed a substantial aggravation of colitis phenotype when adeno-DPP4 injected mice were further treated with a low-dose of DSS in comparing the mice that were kept on only a low-dose of DSS or just adeno-DPP4 injected mice (**Fig. 2A-F**). These data imply that DPP4 contributes to an amplification of colitogenic insults leading to aggravated pathological inflammation.

**Figure 2.**
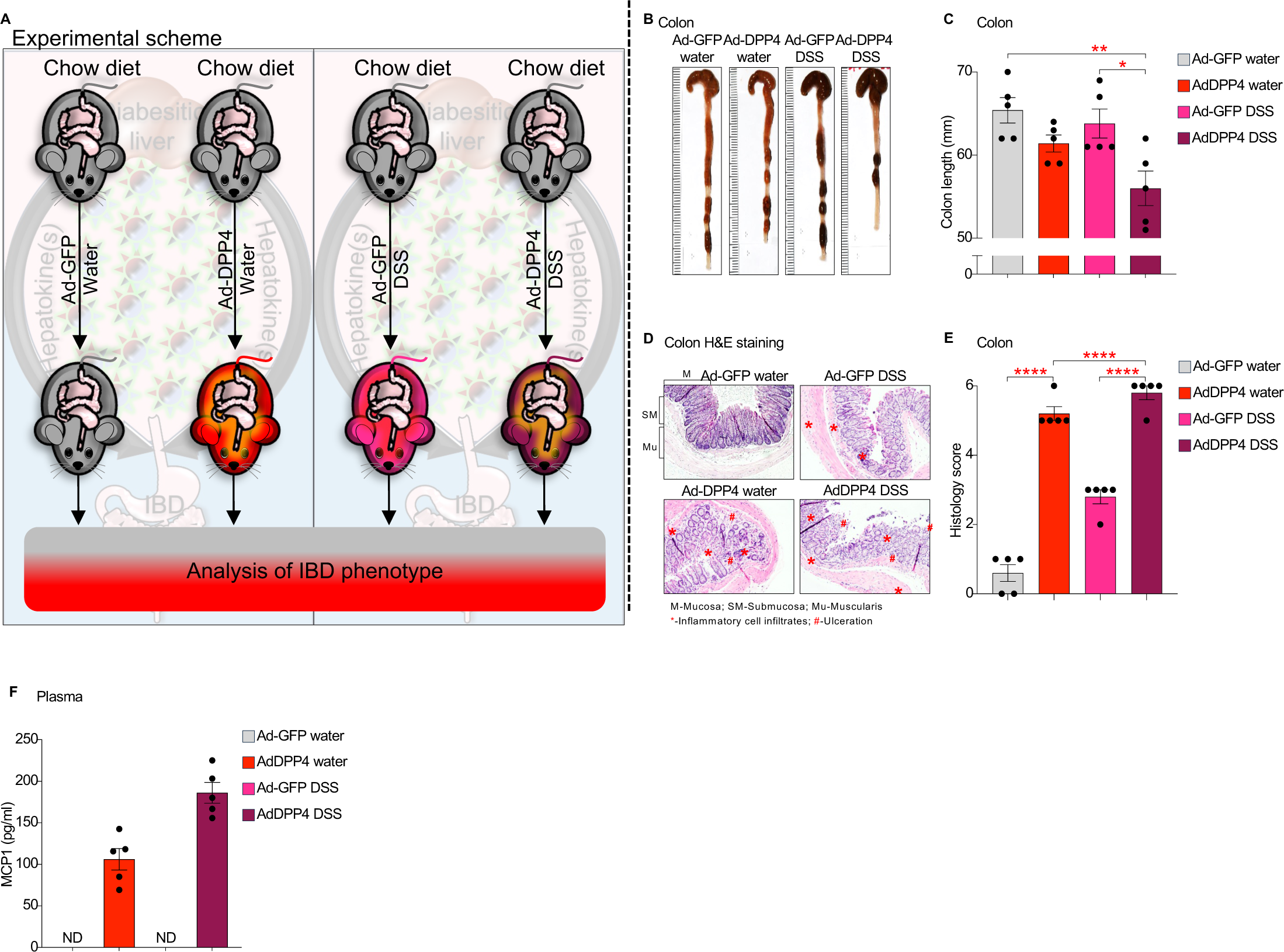
Overexpression of DPP4 augments DSS-induced colitis. **A**, Experimental design. WT chow-fed mice were injected with adenovirus expressing GFP (Ad-GFP) or Adenovirus over expressing DPP4 (Ad-DPP4) for 7 days, followed which they were administered DSS in drinking water. **B & C**, Representation of colon with measurement of colon length. **D & E**, H&E staining representation of colon sections with histopathological scoring. **F**, Circulatory plasma MCP1 levels in mice. N = 5 mice per group; Data are mean ± SEM; ns, non-significant; *, < 0.05; **, < 0.01; ***, < 0.001; ****, < 0.0001 by one way ANOVA. ND = Not determined.

### Hepatocyte–DPP4 is required to aggravate colon inflammation in diabesitic mice

Disruption of calcium homeostasis in the diabetic liver results in excessive accumulation of cytosolic calcium that triggers altered liver pathophysiology(19). One of the net effects is the overstimulation of calcium-dependent CAMKII that relays ATF4-signal transduction to produce DPP4 by the diabetic liver(7). To substantiate the contribution of liver secretory DPP4 in instigating diabetes-linked-colon inflammation, we knocked down DPP4 levels specifically in the livers of HFD-fed prediabetic mice using adeno-associated viruses (type 8) expressing anti-*Dpp4* shRNA. Post-four weeks of AAV8-shDpp4 injections, we analyzed DPP4 expression in 7 different tissues and found DPP4 expression is reduced to 90-95% only in liver. The expression of DPP4 is unaffected in other tissues confirming liver-specific knock down of DPP4. Additionally, liver silencing of DPP4 lowered circulatory DPP4 activity by 55%-60% suggesting a dominant contribution of liver secretory DPP4 (**Supplementary Fig. S5A-O**). WT vs liver-specific DPP4 silenced mice were used to investigate the contribution of liver-secretory DPP4 in colitis post-additional colitogenic stimuli of low-dose DSS (**Fig. 3A**). Silencing of Liver-DPP4 resulted in striking resistance to decrease in colon length (shScr DSS vs shDpp4 DSS), reduced colon pathology, and retained mucus secretion ability of colons (**Fig. 3B-I**). We also found that inflammatory markers relevant to human colitis, *Mcp1, Tnfa, Il-8, Il-1b, Il-6, Lcn2, Crp, Il-1ra,* and *Tff3* are largely reduced upon liver-specific silencing of DPP4 (**Fig. 3J-R**). Additional assays to determine gut barrier functionality confirmed liver DPP4’s contribution to compromised gut integrity in diabetic mice (**Fig. 3S-V**). As alluded to above, diabetes is a disease with multiple comorbidities. Weakening of the immune system makes diabetic patients vulnerable to infections(20). In the context of diabetic colitis, we evaluated acquired enteric infections as leading colitogenic stimuli. We established diabetic salmonellosis mice cohorts by oral gavaging of *Salmonella enterica* serovar Typhimurium (STM). Mice that are infected with oral dosing of STM showed rapid loss of BW and splenomegaly without alteration of VAT and liver mass (**Supplementary Fig. S6A-D**). To validate the role of liver-DPP4 in enteric infections, we set up a cohort of WT and liver-silenced-DPP4 mice that were either subjected to STM infection or left uninfected. We observed that liver-specific lowering of DPP4 protected mice from STM-induced colonic pathological inflammation (**Fig. 4A-K**). Further, mice with liver-silenced DPP4 provided benefits like higher mucin secretion (**Fig. 4L-O**), and maintenance of gut integrity (**Fig. 4P-S**) to maintain a healthy gut to provide protection from STM-triggered salmonellosis.

**Figure 3.**
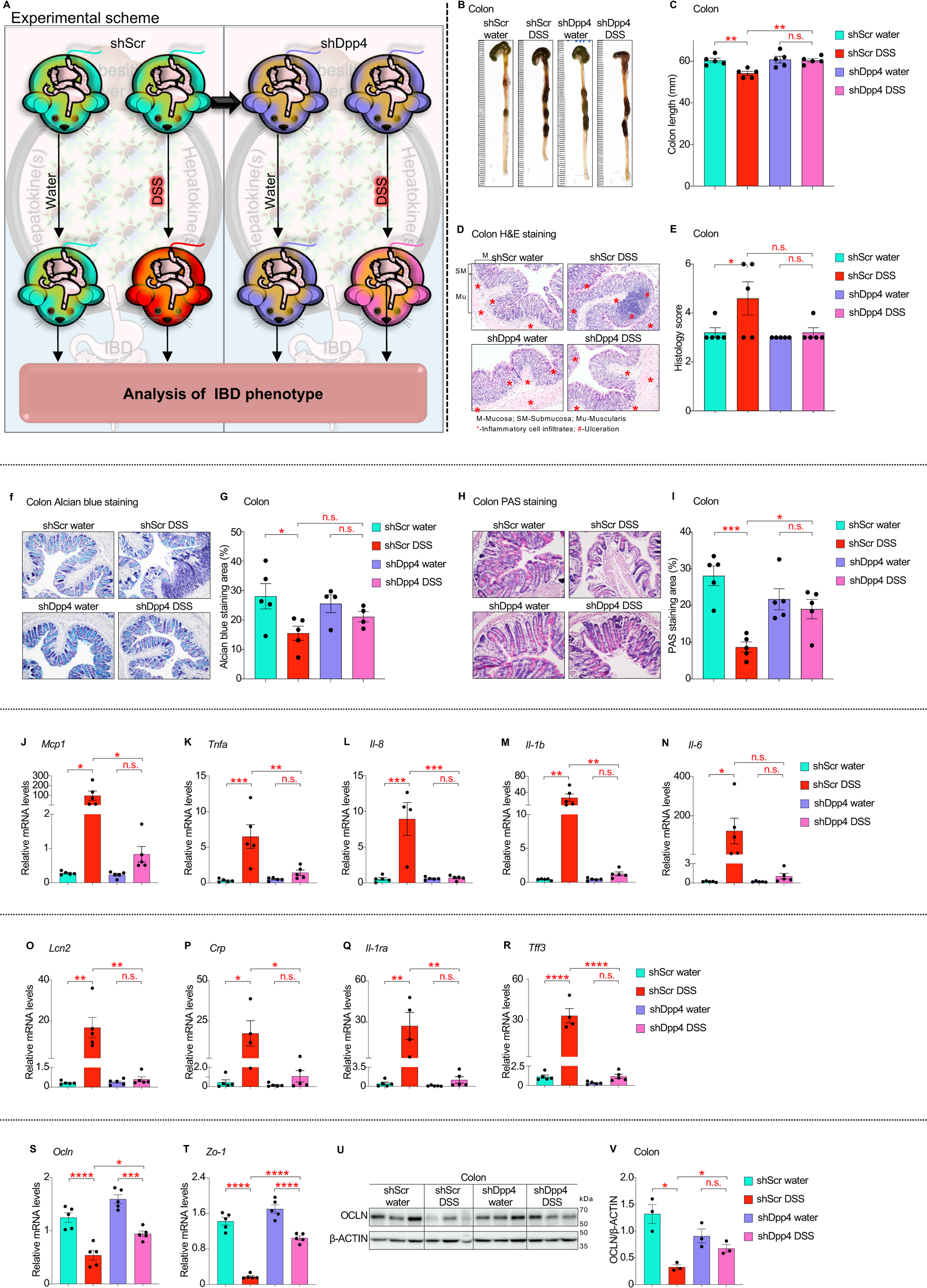
Hepatocyte–specific DPP4 silencing alleviates colon inflammation in DSS-challenged mice. **A.** HFD-fed mice were intravenously injected with AAV8-shDPP4 viruses (ShDpp4) and with control AAV8-shScr viruses (shScr). After 4 weeks post injections mice were challenged with colitogenic stimuli, dextran sodium sulfate (DSS) for 5 days and were analysed as follows: **B & C**, Representative colon images with quantified colon length. **D & E**, Histopathological analysis of colon tissues by H&E staining and histological scoring. **F & G**. Acidic mucin representation in the colon by Alcian blue staining with quantification. **H & I**, Neutral mucin quantification in colon tissue by PAS staining along with stain area quantification. **J-T**, *Mcp1, Tnfa, Il-8, Il-1b, Il-6, Lcn2, Crp, Il-ra, Tff3, Ocln,* and *Zo-1* mRNA levels in colon tissue. **U & V**, Colonic expression levels of OCLN with relative quantification by densitometry. N = 3-5 mice per group; Data are mean ± SEM; ns, non-significant; *, < 0.05; **, < 0.01; ***, < 0.001; ****, < 0.0001 by one way ANOVA.

**Figure 4.**
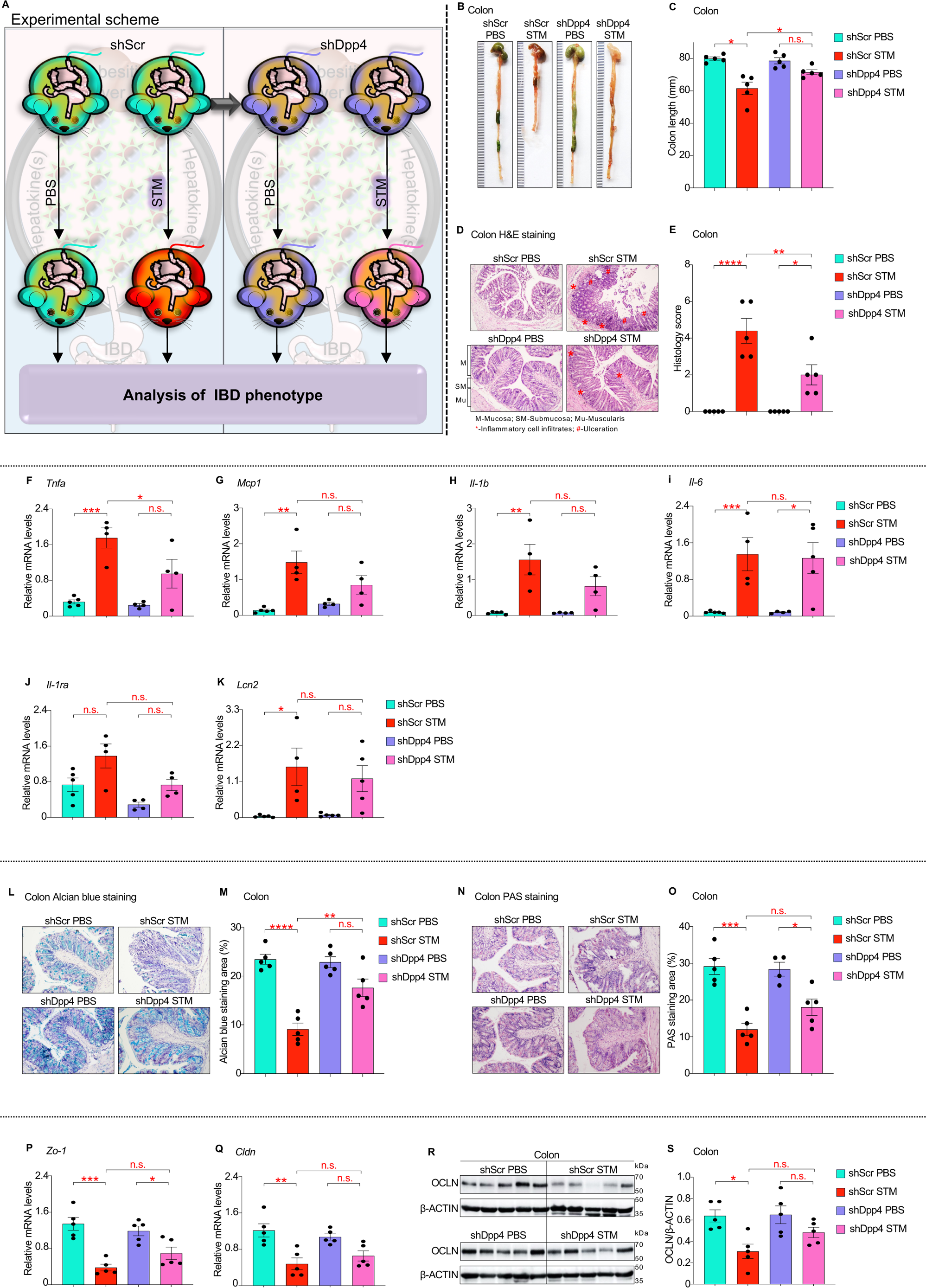
Liver secretory DPP4 elevates *Salmonella enterica* serovar Typhimurium (STM) infected colon inflammation. **A.** Liver-specific liver silencing was achieved as described in Figure 3. shScr injected control cohorts and shDpp4 injected test cohorts were then infected with STM peroral and the following analyses were performed. **B-I**, Colon pathology and inflammation analyses by measuring colon length (**B & C**), Histopathological scoring by H&E staining (**D & E**), and gut inflammatory genes assessment by qRT-PCR analysis of *Tnfa, Mcp1, Il-1b, Il-6, Il-1ea, Lcn2* (**F - K**). Gut mucin production estimation by Alcian blue and PAS staining (**L-O**) and gut permeability assessment by *Zo-1*, *Cldn* transcript levels (**P & Q**), and protein levels of tight junction protein OCLN using western blotting (**R & S**). N = 5 mice per group; Data are mean ± SEM; ns, non-significant; *, < 0.05; **, < 0.01; ***, < 0.001; ****, < 0.0001 by one way ANOVA.

### Silencing of hepatocyte DPP4 levels, but not systemic inhibition of DPP4 activity by sitagliptin mitigates DSS-challenged colon inflammation

DPP4 is an N-terminal dipeptidase that inactivates its target substrates by cleaving off two amino acids from the N-terminal of the peptide(21). One of the classical cognate substrates of DPP4 is glucagon-like peptide-2 (GLP2)(22), an intestinotrophic hormone that promotes gut homeostasis via promoting enteric cell proliferation, inhibition of apoptosis, maintenance of mucosal integrity and reduction of gut permeability(23–25). Thus, we hypothesize that in the setting of diabetes elevated levels of liver-DPP4 target GLP2 thereby lowering gut protection and promoting pathological inflammation. We utilized a clinically approved DPP4 activity inhibitor sitagliptin to test this hypothesis. We performed simultaneous experiments with liver-specific silencing of DPP4 and systemic inhibition of DPP4 activity in a prediabetic HFD-fed mouse cohort. Treatment of mice with AAV8-shDpp4 injections lowered only liver DPP4 levels without affecting colonic DPP4 expression, while administration of sita had no effect on DPP4 protein levels in liver and colon tissues (**Supplementary Fig. S7A-D**). In consistent to the above notion, we found that knockdown of liver DPP4 resisted a decrease in DSS-instigated colon length with a lowering of colon pathology score (**Fig. 5A-D**). Despite sitagliptin lowered DPP4 activity (**Supplementary Fig. S7E**) it could not protect mice from DSS-triggered colon pathology (**Fig. 5A-D**). Similarly, at the molecular level liver-specific silencing of DPP4, but not the inhibition of DPP4 activity altered the course of colonic production of mucin, colon inflammation, and gut barrier dysfunction (**Fig. 5E-Q**). Although sitagliptin lowers DPP4 activity, it compensates by over-production of circulatory DPP4 protein (**Supplementary Fig. S7E, F**). Our this observation is consistent with the report that DPP4 is known to instigate receptor-mediated inflammatory pathways in adipose tissue macrophages(7,26). These combined data are supportive of the idea that DPP4 independent of its enzymatic activity, acts as a ligand to initiate gut inflammatory pathways.

**Figure 5.**
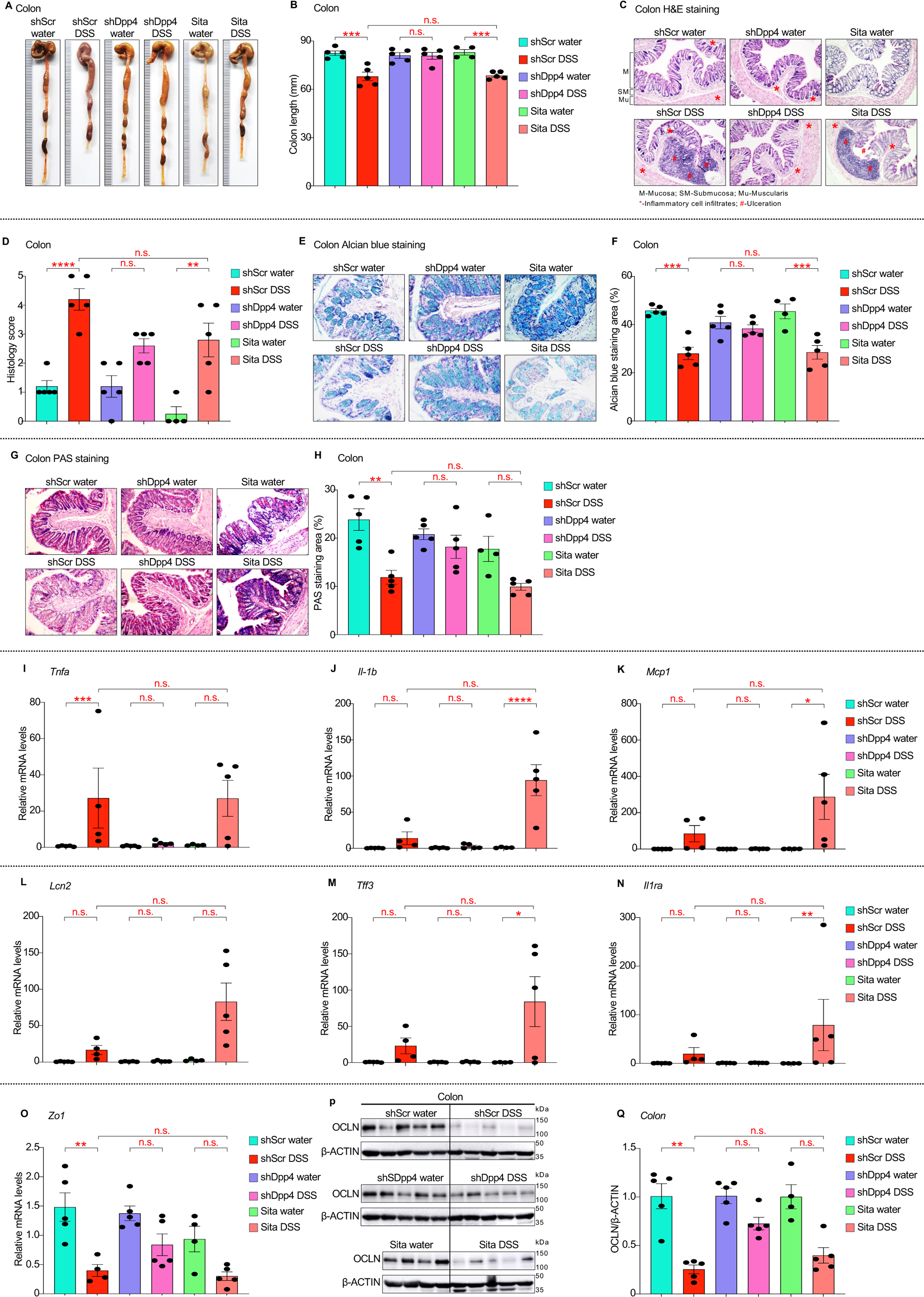
Silencing of hepatocyte DPP4 levels, but not systemic inhibition of DPP4 activity by sitagliptin mitigates DSS-challenged colon inflammation. Liver-specific silencing of DPP4 was achieved as described in Figure 3. Mice were administered with 0.3mg/ml of sitagliptin to achieve systemic inhibition of DPP4 activity and then challenged with colitogenic DSS stimulation. Control (shScr), Liver-specific DPP4 silenced (shDpp4), and systemic DPP4 activity inhibition (Sita-treated) mice were analysed as follows: **A & B**, Colon images and colon length measurement. **C & D**, H & E-stained colon tissues with respective quantifications. **E & F**, Alcian blue staining of colon tissues along with quantification of acidic mucin area of the colon. **G & H**, PAS staining for neutral mucin quantification**. I-O**, *Tnfa, IL-1b, Mcp1, Lcn2, Tff3, Il-1ra,* and *Zo-1* mRNA levels in colons. **P & Q** protein levels of tight junction protein OCLN using western blotting. N = 5 mice per group; Data are mean ± SEM; ns, non-significant; *, < 0.05; **, < 0.01; ***, < 0.001; ****, < 0.0001 by one way ANOVA.

### Liver-secretory DPP4 suppresses the autophagic pathway to flare gut inflammation

To understand the mechanistic details of DPP4-mediated aggravation of gut inflammation in diabetes, we performed an unbiased proteomics analysis of colon tissues obtained from the control and liver-silenced DPP4 cohort. Principal component analysis scores indicated a differential expression of proteins in these cohorts (**Fig. 6A**). Detailed protein expression analysis of DSS-treated (shScr DSS vs shDpp4 DSS) cohorts was performed after data normalization with protein expressions in respective water-treated mice cohorts. A total of 624 proteins were upregulated in shDpp4 cohort while 143 genes were upregulated in shScr DSS cohort (**Fig. 6B**). Gene ontology pathway analysis highlighted “Autophagic pathways” as a top-upregulated pathway (**Fig. 6C**). A cohort of literature suggests autophagy’s positive impact on maintaining gut health(27–29). Defects in autophagic response have been suggested to be at the core of pathological inflammation in the gut(16,30,31). From a clinical point, autophagy inducers are in clinical trials for treating inflammatory bowel disease(32–35). Thus, we propose, during diabesity liver DPP4 targets gut autophagy to aggravate gut inflammation. In support of our thought, we curated mass-spectrometry data and identified striking induction of autophagic proteins upon liver-specific DPP4 silencing (**Fig. 6D**). This data is further verified by RT-qPCR analysis of autophagy responder gene’s *Atg5, Ulk1, Atg7, Nrf1, Atf7,* and *Lamp1* expressions (**Fig. 6E-J**). We also identified that LC3B levels were markedly reduced in colon samples obtained from control shScr-DSS treated mice. In contrast, liver-specific silencing of DPP4 enhanced LC3B expression in shDpp4-treated colons even after treatment with experimental DSS insults (**Fig. 6K-L**). Collectively, our data suggest a crucial role of liver DPP4 in instigating gut inflammation via the inhibition of autophagy leading to increased vulnerability of gut to diabesity-associated pathological inflammation.

**Figure 6.**
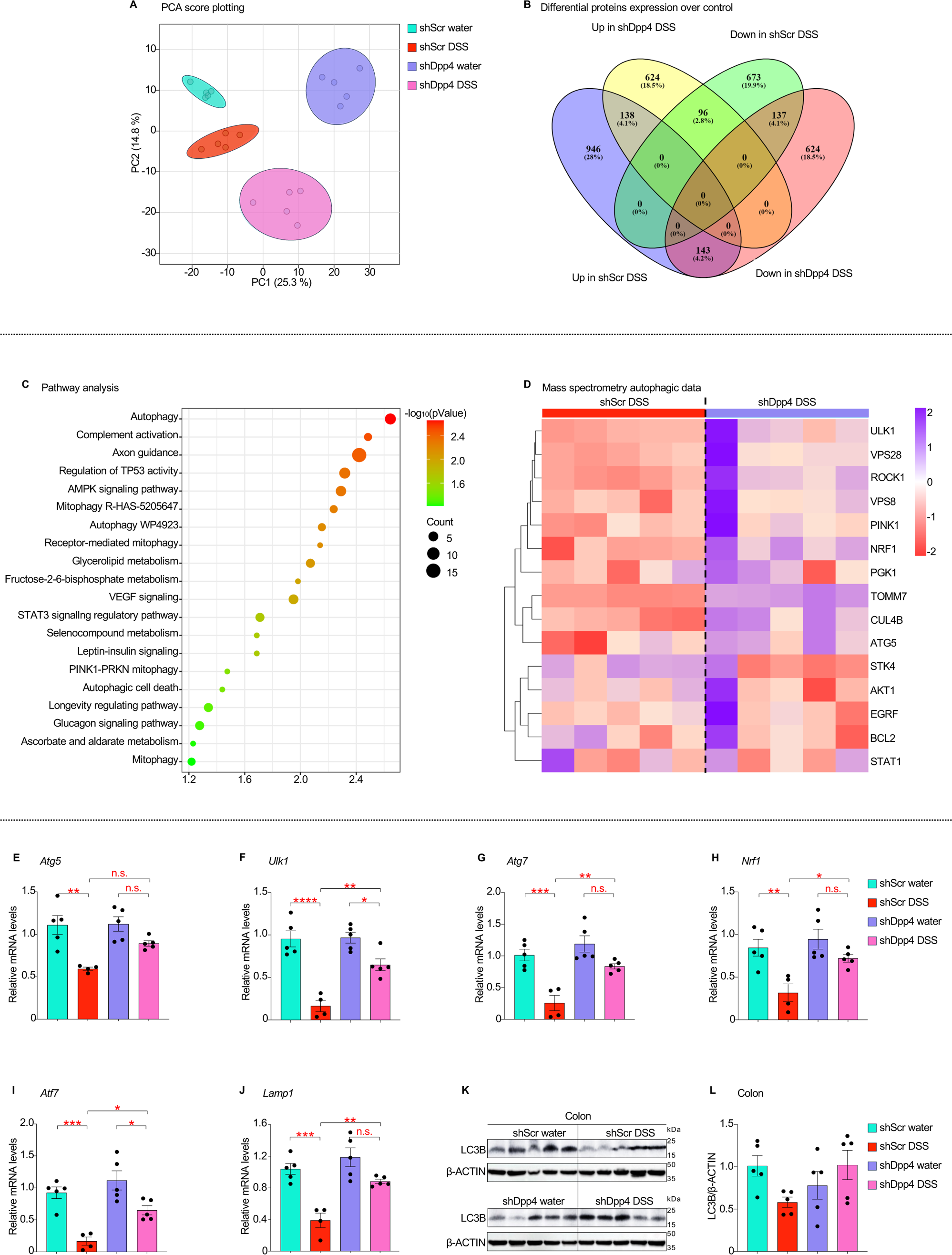
Liver-secretory DPP4 suppresses the autophagic pathway to flare gut inflammation. The total lysate of colon tissues obtained from four cohorts of mice from Figure 3 were analysed by unbiased proteomics and following differential analyses were performed on mass spectrometry data. **(A)** Principal component analysis (PCA) of four cohorts of mice. **(B)** After adjusting for protein expressions with respective control mice (shScr DSS over shScr water and shDpp4 DSS over shDpp4 water) differential protein expressions were analysed. **(C)** Gene ontology pathway analysis. **(D)** Heat map for autophagic protein expressions. **E-L**, mRNA expressions of *Atg5, Ulk1, Atg7, Nrf1, Atf7,*and *lamp1* were analysed by qRT-PCR and protein levels of LC3B were estimated by western blotting. LC3B was quantified by densitometric analysis and presented over β-ACTIN. N = 5 mice per group; Data are mean ± SEM; ns, non-significant; *, < 0.05; **, < 0.01; ***, < 0.001; ****, < 0.0001 by one way ANOVA.

**Figure 7.**
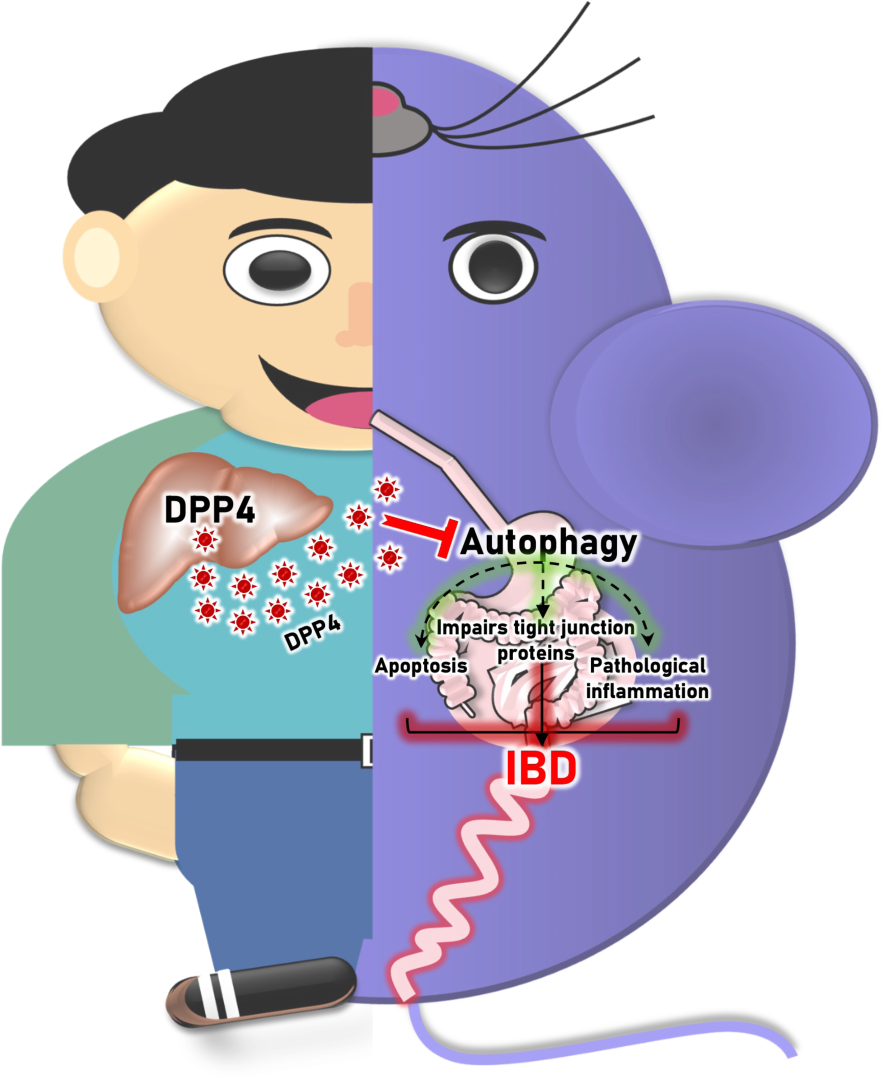
Summary scheme: Data obtained from liver and plasma of NASH, NAFLD and DM humans and mice cohorts of high-fat diet-induced and genetic diabesitic models (HFD and Db/Db mice) suggest that secretory DPP4 is enriched in human and mice livers. The secretory DPP4 attenuates gut autophagic pathway which is at the core of high apoptosis of enterocytes, impaired gut permeability, and pathological inflammation of diabetic gut.

## Discussion

Diabesity affects energy homeostasis having broader impairment effects on multiorgan function. Of note a parallel rise in cases of diabesity and IBD are of major concern with low or limited knowledge of causation(36). Liver secretory hepatokines serve as mediators of cross-communication to a variety of organs like VAT, pancreas, muscle etc. affecting the physiological and pathological output. For example, using the mouse models of prediabetes the role of hepatocyte secretory DPP4 in mediating adipose tissue macrophage (ATM) inflammation that leads to glucose intolerance and insulin resistance has been illustrated(7). However, the role of hepatokines in inflaming the gut is not well illustrated and this becomes extremely important considering the high prevalence of IBD cases in diabetic patients. Using the combination of data obtained from livers, blood samples of human diabetes and related comorbidities and gain and/or loss of functions in mouse models of pre-diabetes/morbid diabetes we provide evidence for diabetic liver enriched DPP4 to fuel gut inflammation via suppressing protective autophagic response.

Colitis is marked by increased death of enteric cells with a concomitant decrease in epithelial cell proliferation, infiltration of immune cells to the site of damage, and accumulation of inflammatory milieu(14). These classical signs of colitis are evident in diabetes patients with unknown etiology. Considering the liver endocrine functions are elevated in diabetes and the potential of hepatokines in inflammatory diabetes-associated comorbidities including colitis, a data curation analysis performed on the liver, and blood samples obtained from seven different cohorts of human metabolic disorders like NASH (4 human cohorts), NAFLD (1 human cohort), and diabetes mellitus (2 human cohorts) demonstrating DPP4 enrichment in the livers and blood samples obtained from all human cohorts. Our recapitulation data using prediabetes and genetically morbid obese-diabetic db/db mice confirms these human findings. Interestingly, liver-enriched DPP4 levels are positively associated with colitis markers in these murine models implying the possible role of liver DPP4 in the occurrence of diabetes-linked colitis. We further supported this idea by using the gain of function and hepatic loss of DPP4 levels in mouse models. We report that the overexpression and/or knockdown of liver secretory DPP4 levels directly influences gut pathological inflammation. For example, DPP4 overexpression in non-diabetic mice alone is able to induce colitis and combinatorial DPP4 overexpression with DSS treatment aggravates experimental colitis. On the contrary, hepatic silencing of DPP4 in diabetic mice provides protection from DSS-induced colitogenic insults. Recent findings suggest that IBD patients with diabetic pathology are prone to encounter a variety of infections including enteric colonization of opportunistic pathogens(20). *Salmonella* is known to establish enteric infections by degrading the mucin layer and breaching the gut barrier. Reducing liver DPP4 levels with AAV8-shDPP4 injections leads to a significant reduction in *Salmonella*-mediated gut pathology suggesting a beneficial role of liver DPP4 silencing in colitis.

DPP4 is one of the exceptional hepatokines, that exhibits dual functionality, one is peptidase activity, and another is the engagement of cell surface receptors to trigger inflammatory pathways(37). Utilizing peptidase ability DPP4 inactivates its target substrates by cleaving off two N-terminal amino acids. One of the gut-health-promoting hormones, GLP2 is a well-known substrate-target of DPP4. GLP2 is known to promote gut health via engaging with GLP2 receptors on enteric cells(25). GLP2-GLP2R engagement transmits a signal to promote epithelial cell proliferation, inhibits apoptosis, enhances mucin production and facilitates gut barrier functions(24,25). The expression of DPP4 is enhanced under diabetic conditions suggesting the impairment of GLP2 functionality(38). To test DPP4 targeted GLP2 axis in promoting gut inflammation, we inhibited DPP4 enzymatic activity by peroral administration of sitagliptin, a clinically approved DPP4 drug to enhance glucose tolerability. Although sitagliptin inhibited DPP4 enzymatic activity, it had no positive impact on gut inflammation in the experimental colitis model. Detailed investigation revealed that although sitagliptin inhibited DPP4 activity, it enhanced the production of DPP4 protein. This data is consistent with previous findings and suggests that DPP4 promotes gut inflammation via interacting with its inflammatory receptors in the gut. A study by Moran et al. identified lower serum DPP4 levels in active Crohn’s disease patients (n=23) compared to healthy controls (n=17) implying a compensatory mechanism to counter DPP4-instigated gut pathological inflammation(39). A large population-based longitudinal study has clearly stated about 75% increase in the risk of IBD in type 2 diabetic patients that are on a DPP4-inhibitor regime and advocates a word of caution to prescribe sitagliptin to type 2 diabetic patients as it may be an independent risk factor for the diabetes-associated prevalence of gastric problems(40).

To unravel a connecting link between diabetic-liver enriched DPP4 and colonic pathological inflammation, we performed an unbiased proteomics analysis. Pathway enrichment analysis suggests “autophagy” is the top pathway upregulated in the liver DPP4 silenced cohort. This observation is a corroboration of recent findings. A group of studies have recently reported that increased autophagic response counters pathological inflammation in the gut by clearance of damage-associated molecular signals, regulation of immune response, maintenance of gut integrity and mucus secretion(29). Contrary defective autophagy leads to aggravated colon inflammation(27). Liver-specific silencing of DPP4 induces the autophagic gene profile in DSS-treated experimental colitis with significant activation of LC3B, a cardinal feature of autophagy. These data indicate the vital role of diabetic liver DPP4 in the impairment of gut autophagy and aggravated colon inflammation. In conclusion, there is mixed literature about the role of DPP4 and its inhibitors in providing protection or aggravation of colitis. One of the major low sides of these studies is a lack of mechanistic knowledge of whether DPP4 activity is crucial or DPP4-mediated inflammatory signaling in vital for colonic pathological inflammation. Our discovery highlights the “liver-DPP4-autophagy inhibition axis” in pathological colon inflammation and provides an opportunity to target this nexus to combat diabesity-linked comorbid IBD.

## Methods

### Mouse experiments

All animal studies were performed following the guidelines established by the Institute Animal Ethics Committee (IAEC) on Use and Care of Animals at the National Institute of Immunology. Mice were housed in the specific pathogen-free facility under 12/12 h light/dark cycles with free access to food and water. All animals were acclimatized for one week in the animal facility before commencement of experiments. For chow feeding, mice were fed with regular chow diet. For HFD feeding, mice were fed with a diet containing 60 kcal % fat. For establishment of diabesity phenotype in mice, male C57BL/6J mice were fed a high-fat, high-calorie diet (HFD, 60 kcal % from fat) for 16 weeks starting at six weeks of age and were maintained on a 12-h light–dark cycle. Six-week-old chow-fed male Db/Db mice were used as genetically diabesitic mice model. Intraperitoneal-glucose tolerance tests were performed by intraperitoneal injection of glucose (1 g/kg body weight for DIO, 0.5 g/kg body weight for Db/Db mice) following an overnight fast. To obtain DPP4 overexpression mice, six-week-old lean male C57BL/6J mice were injected via tail vein with Ad-DPP4 virus and Ad-GFP as control. Similarly, to obtain hepatocyte specific DPP4 silenced mice, eight-week-old wild-type DIO mice were injected with AAV8–H1-shDPP4 or AAV8–H1-scramble (shScr as control). In all mouse experiments, recombinant adenovirus (1 × 10^9^ plaque-forming units per mouse) or adeno-associated virus (1 × 10^15^ genome copies per mouse) was delivered by tail vein injection, and experiments were commenced after 10 days for the experiment in Fig. 2, or after 4 weeks for the experiments in Fig.3, 4 and 5. DIO mice were treated for four weeks with sitagliptin by adding it to the drinking water at 0.3 mg/ml, which results in a dose of ∼30–45 mg/kg/day. Mouse plasma samples were collected from lean or obese mice after 4–5h of food withdrawal, with free access to water. Blood glucose was measured after 4–5h of food withdrawal in mice using a glucose meter (ACCU-CHECK instant, ROCHE diagnostics). At a predetermined time, mice were euthanized according to relevant guidelines, and the wet weights of tissues such as the liver, VAT, and spleen were measured. Colon length was measured immediately after harvesting, and photographs were taken before the colon was flushed with ice-cold PBS. Samples intended for qPCR, protein analysis, or further study were snap-frozen in liquid nitrogen immediately. Distal part of the fresh cleaned colon tissue was fixed in PBS-buffered 10% formalin and used for histological analysis. Additionally, mice were handled with extra care to minimize stress during blood glucose measurement and blood collection.

We did not predetermine the sample size; instead, we used group sizes that are typical for this type of study in the literature. Mice of the same genotype were randomly assigned to experimental or control groups to minimize potential bias. For instance, age and weight - matched WT and liver-silenced DPP4 mice were randomly allocated to either water or DSS treatment. The investigators were not blinded to the group allocations during the experiments and outcome assessments. Mice were excluded from the study if they exhibited skin lesions from fighting, growth retardation resulting in weight loss of 10% of their initial body weight, or showed signs of illness requiring euthanasia. Based on these criteria, typically between zero and/or one mouse was removed before analysis.

### Experimental colitis induction in mice

For the experiments shown in Figure 2, 8-week-old mice of the specified genotypes were used, while 12-week-old mice were utilized for the experiments in Figures 3, 4, and 5. To induce an acute colitis phenotype, the mice were given either 1% or 2% dextran Sulfate sodium (DSS) in their autoclaved drinking water for 5 days. In the dose kinetics experiment (Figure S4), DSS concentrations ranging from 1% to 5% were tested to determine the optimal dose for the primary experiments. Enteropathogenic colitis was induced by administering a single dose of 10^6^ CFU of *Salmonella enterica* serovar Typhimurium (STM). Mice were sacrificed 4 days post-infection for analysis, and body weight changes in *Salmonella*-infected mice were monitored over time.

### Microarray data acquisition

The Gene Expression Omnibus (GEO) database is a publicly accessible repository for high-throughput gene sequencing data. We selected datasets based on two criteria: (A) liver or blood tissue samples were clinically diagnosed with metabolic syndromes such as NAFLD, NASH, or diabesity, and (B) normal liver tissues or blood samples from apparently healthy participants were used as negative controls. In this study, we selected seven datasets (GSE24807, GSE17470, GSE63067, GSE135251, GSE167523, GSE26168, GSE250283) from the GEO database. The characteristics of these datasets are detailed in the main text. GEO2R, an R-based tool available on the GEO website, was used to identify differentially expressed genes (DEGs) between groups. For our analysis using GEO2R, we focused on 12 established hepatokines associated with metabolic syndromes and compared them between diseased and healthy participants. We applied a cutoff of log fold change >0.5 and an adjusted p-value <0.05, using the Benjamini & Hochberg method for p-value adjustment. Subsequently, we used GraphPad software to plot the quantitative data obtained from the datasets.

### Quantitative reverse transcription–PCR (qRT-PCR) analysis

Total RNA was extracted from tissue samples of multiple organs using TRIzol method. The concentration, purity and quality of the isolated RNA was measured using Nanophotometer N120 (Implen). 2μg total RNA was reverse transcribed using 1^st^ strand cDNA synthesis kit. Real time quantitative (q) PCR was performed with CFX opus 384 and CFX opus 96 (Biorad) using TB Green Premix Ex taq and gene-specific primers. The relative expression levels were normalized by the expression levels of 18S or GAPDH. The sequences of gene-specific primers used for the RT-qPCR assays are enlisted in Supplemental Table 2.

### Immunoblotting

Tissue samples from multiple organs were homogenized in RIPA lysis buffer supplemented with protease phosphatase inhibitor. Lysates were cleared by centrifugation using centrifuge 5424R (Eppendorf) at 15,000 rpm for 15 min at 4 °C. The protein content of lysates was measured using the DC Protein Assay kit with a Clariostar plate reader from BMG Labtech. Equal amount of protein from tissue was resolved on 10-15% SDS-PAGE gels and then transferred to PVDF membranes. The membranes were blocked at room temperature for 1 hour using 5% non-fat milk followed by incubation with primary antibodies overnight. Blots were then washed thoroughly with TBST and probed with HRP-conjugated secondary antibodies for 1 hour. Protein bands were detected using the Clarity Western ECL substrate Bio-Rad and imaged with the Image Quant 500 system (Amersham). To ensure equal protein loading, the same blot was stripped with stripping buffer (1.5% glycine + 0.1% SDS, Tween20 1ml, pH 2.2) and then incubated with an HRP-conjugated anti–mouse GAPDH or β-actin antibody. Relative intensities of protein bands were quantified using ImageJ analysis software.

### Histological analysis of murine colon tissues

For histological analysis, we removed the colon from mice and distal part of colon was fixed in 10% formalin. The samples were embedded in paraffin and cut into 5 µm sections. After rehydration and permeabilization, we stained the sections using haematoxylin and eosin (H&E), alcian blue, and Periodic Acid-Schiff (PAS). To study the histopathological characteristics of DSS- and water-treated mice, the slides were initially stained with haematoxylin for 7 minutes, rinsed in tap water, and then immersed in acid alcohol to minimize nonspecific staining. Bluing was conducted in a 0.1% sodium bicarbonate solution to enhance the stain color. Subsequently, the slides were counterstained with eosin by dipping them 30 times in the stain. The sections were then dehydrated, cleared in xylene, and mounted with DPX for imaging. For pathological assessment, the H&E-stained sections were evaluated using light microscopy following the criteria of histological score previously reported(41). The observer scored the tissue morphology as follows: The extent of the leukocyte infiltration was scored as 1, significant increase in mucosa (mild); 2, confluence in the submucosal part(moderate); 3, transmural infiltration (marked). The epithelial damage was scored as 1, focal erosion; 2, focal ulceration; and 3, extended ulceration and granulomatous tissue. For mucin quantification, slides were stained with alcian blue for 30 minutes, then rinsed and counterstained with haematoxylin for 7 minutes. To minimize nonspecific alcian blue reactions and enhance the blue color of haematoxylin-stained nuclei, the slides were treated with acid alcohol and sodium bicarbonate solution. The sections were subsequently dehydrated, cleared in xylene, and mounted with DPX for imaging. The area stained by alcian blue (%) was quantified using ImageJ software. For PAS stain, the sections were incubated in 1% periodic acid, rinsed in tap water, and then incubated in Schiff’s reagent for 7 minutes. Following this, the slides were washed and counterstained with haematoxylin for 5 minutes. To reduce nonspecific PAS reactions and enhance the blue color of haematoxylin-stained nuclei, the slides were treated with acid alcohol and sodium bicarbonate solution. The area stained by PAS stain (%) was quantified using ImageJ software.

### AAV8-shScr and AAV8-shDpp4 preparation

A short hairpin RNA (shRNA) targeting DPP4 was cloned into an adeno-associated virus 8 (AAV8) expression construct and AAV8-sh-DPP4 expressing viruses were produced in the laboratory by triple transfecting HEK293T cells. The control cohort of mice were injected with AAV8 viruses encoding shScr construct. Briefly, PEI-DNA complex was added to the 70 – 80% confluent HEK293T cells, post 24 hours fresh media was added along with tryptone and sodium butyrate mixture. Post 72 hours cells were scrapped and virus was harvested by PEG precipitation. Virus titer was estimated by measuring viral genome copies using qRT-PCR against ITR.

### DPP4 activity assay

Plasma DPP4 activity was estimated using a fluorometric assay kit as per standard manufacturer protocol (Enzo Life Sciences).

### DPP4 and MCP1 ELISA

DPP4 protein levels were estimated using a DPP4 ELISA kit procured from R&D systems as per standard manufacturer instructions. MCP1 protein levels were estimated using a MCP1 ELISA kit procured from Peprotech as per standard manufacturer instructions.

### Unbiased proteomic analysis in murine colon tissues

Proteins were isolated from the colons of four groups; ShScr water, ShScr DSS, ShDpp4 water, and ShDpp4 DSS (n=5 mice per group). Briefly, liver tissues were lysed using RIPA lysis buffer with sonication (30 sec on, 10 sec off, 3 cycles) and vortexing. Samples were then centrifuged for 15 min at 12,000 g (4°C) twice. Protein precipitation was performed by adding ice-cold acetone up to the brim of the microcentrifuge tube, and samples were incubated at −20°C overnight. The following morning, samples were centrifuged and the pellet was processed for proteomic analysis. The pellet was resuspended in 50 μl ABC buffer and incubated at 4°C for 30 minutes. Protein concentrations were quantified using the Bradford Protein Assay Kit. 50 μg of protein was reduced by adding 20 μl DTT and incubating for 1 hour at 60°C. This was followed by IAA treatment (20 μl) to block reduced cysteine residues, with a 30-minute incubation in darkness. The protein suspensions were digested overnight with 10 μl trypsin at 37°C in a water bath. The next day, 5 μl of formic acid was added to stop trypsinization. The digested peptides were then desalted on C18 columns (Thermo). The columns were activated with 200 μl Buffer A (50% acetonitrile), washed, and equilibrated with 200 μl Buffer B (5% acetonitrile). The samples were loaded and passed through the column four times. The column was then eluted with Buffer C (30 μl, 40 μl, 30 μl) (90% acetonitrile), and the pass-through was centrifuged. Samples were concentrated in a vacuum dryer, and the pellet was resuspended in 30 μl of 2% acetonitrile in 0.1% formic acid. The suspension was centrifuged, and 15 μl of the samples were loaded into the MS vial for analysis. Peptide solutions were subjected for 2-hour LC/MS/MS with Vanquish Neo-UHPLC system interfaced to a Thermo Orbitrap Exploris 240.

### Statistical analysis

All results are presented as mean ± SEM, with statistical significance indicated as ns, non-significant; *, < 0.05; **, < 0.01; ***, < 0.001; ****, < 0.0001. Statistical analyses began with assessing data distribution using the Shapiro-Wilk test. For data with a Gaussian distribution, comparisons between two groups were analyzed using an unpaired Student’s t-test (two-tailed), while non-Gaussian distributed data were analyzed with the Mann-Whitney U test (two-tailed). For comparisons among multiple groups with Gaussian distribution, a one-way ANOVA followed by Tukey’s multiple comparison test was used. For non-Gaussian distributed data, the Kruskal-Wallis test followed by Dunn’s multiple comparison test was employed. Correlation analysis was conducted using Pearson’s test for Gaussian-distributed data or Spearman’s rank test for non-Gaussian data. Statistical analyses were performed using Prism software (GraphPad).

## Supporting information

Resource table

## Author contribution

### Contributions

M.A., R.B., and D.S.G. conceptualized the study, designed the experiments, and analysed data. M.A., R.B., C.G., and P.M. conducted the experiments. P.K. helped with the interpretation of data. M.A., M.Y., and J.S.M. conducted mass spectrometric analysis and helped design these experiments and analyse the data. M.A. and D.S.G. wrote the manuscript All the authors reviewed the manuscripts.

## Funding

This work is supported by research grants obtained from SERB CRG grant (CRG/2021/000147), ICMR grant (5/4/8-28/CD/DSG/2022-NCD-II), DBT Ramalingaswami fellowship (BT/RLF/Rentry/38/2019) to D.S.G. and NII core grant.

## Acknowledgement

We thank Mr. Pankaj Kumar for assistance with animal experiments; Dr. Mohd. Ayub Qadri for *Salmonella enterica* serovar Typhimurium strain. Dr. Medha Rajappa, Dr. Pazhanivel Mohan, Dr. S. Gopalan Sampathkumar, Dr. Aneeshkumar A.G., Dr. Santiswarup Singha, and Dr. P. Nagarajan for discussions.

## Competing interest

The authors declare no competing financial interests.

## Data availability

The data that support the findings of this study are available from the corresponding author upon reasonable request.

## Notes

### Competing Interest Statement

The authors have declared no competing interest.

