## Supplementary material for "Diabetic liver-enriched secretory dipeptidyl peptidase 4 (DPP4) fuels gut inflammation via attenuation of autophagy": Resource table

**Supplemental Table 2: qRT-PCR genes specific primers.**

| | | |
|---|---|---|
| Mouse *Dpp4* | Forward | ACCGTGGAAGGTTCTTCTGG |
| | Reverse | CACAAAGAGTAGGACTTGACCC |
| Mouse *Mcp1* | Forward | TTAAAAACCTGGATCGGAACCAA |
| | Reverse | GCATTAGCTTCAGATTTACGGGT |
| Mouse *Il 6* | Forward | AGTTGCCTTCTTGGGACTGA |
| | Reverse | CAGAATTGCCATTGCACAAC |
| Mouse *Lcn2* | Forward | TGGCCCTGAGTGTCATGTG |
| | Reverse | CTCTTGTAGCTCATAGATGGTGC |
| Mouse *Tff3* | Forward | TCTGGCTAATGCTGTTGGTG |
| | Reverse | CTCCTGCAGAGGTTTGAAGC |
| Mouse *Camp* | Forward | GCTGTGGCGGTCACTATCAC |
| | Reverse | TGTCTAGGGACTGCTGGTTGA |
| Mouse *Il1rn* | Forward | GCTCATTGCTGGGTACTTACAA |
| | Reverse | CCAGACTTGGCACAAGACAGG |
| Mouse *Il1ra* | Forward | GGGATACTAACCAGAAGACC |
| | Reverse | GACAGGCACAGCTTGCCCCC |
| Mouse *Pdgfb* | Forward | CATCCGCTCCTTTGATGATCTT |
| | Reverse | GTGCTCGGGTCATGTTCAAGT |
| Mouse *Crp* | Forward | ATGGAGAAGCTACTCTGGTGC |
| | Reverse | ACACACAGTAAAGGTGTTCAGTG |
| Mouse *Tnfa* | Forward | TATGGCTCAGGGTCCAACTC |
| | Reverse | CTCCCTTTGCAGAACTCAGG |
| Mouse *Il1b* | Forward | GCCCATCCTCTGTGACTCAT |
| | Reverse | AGGCCACAGGTATTTTGTCG |
| Mouse *Zo1* | Forward | TTTTTGACAGGGGGAGTGG |
| | Reverse | TGCTGCAGAGGTCAAAGTTCAAG |
| Mouse *Cldn* | Forward | GGGGACAACATCGTGACCG |
| | Reverse | AGGAGTCGAAGACTTTGCACT |
| Mouse *Ocln* | Forward | ATGTCCGGCCGATGCTCTC |
| | Reverse | TTTGGCTGCTCTTGGGTCTGTAT |
| Mouse *Atg5* | Forward | TGTGCTTCGAGATGTGTGGTT |
| | Reverse | GTCAAATAGCTGACTCTTGGCAA |
| Mouse *Ulk1* | Forward | AAGTTCGAGTTCTCTCGCAAG |
| | Reverse | CGATGTTTTCGTGCTTTAGTTCC |
| Mouse *Atg7* | Forward | GTTCGCCCCCTTTAATAGTGC |
| | Reverse | TGAACTCCAACGTCAAGCGG |
| Mouse *Nrf1* | Forward | AGCACGGAGTGACCCAAAC |
| | Reverse | TGTACGTGGCTACATGGACCT |
| Mouse *Atf7* | Forward | ATGGGAGACGACAGACCGTT |
| | Reverse | GGCGTTTGATCTGCAATGATGA |
| Mouse *Lamp1* | Forward | CAGCACTCTTTGAGGTGAAAAAC |
| | Reverse | ACGATCTGAGAACCATTCGCA |
| Mouse *18S* | Forward | CTCAACACGGGAAACCTCAC |
| | Reverse | CGCTCCACCAACTAAGAACG |
| Mouse *Gapdh* | Forward | AGGTCGGTGTGAACGGATTTG |
| | Reverse | TGTAGACCATGTAGTTGAGGTCA |

Supplemental Table 3: Reagents details with catalogue numbers.

| **Reagent** | **Make** | **Catalogue number** |
|---|---|---|
| High fat diet | Research diets Inc, USA | D12492 |

| | | |
|---|---|---|
| Glucose | Sigma | G7021 |
| Injection syringe | BD Glide | 324902 |
| Sitagliptin | Apex Bio | A4036 |
| Ripa lysis buffer | Gbiosciences | 786-490 |
| DC protein assay kit | Biorad | #5000114 |
| PVDF membrane | Merck | 1PVH00010 |
| Skim milk | Millipore | 70166 |
| ECL reagent | Biorad | 1705061 |
| Trizol | Gbiosciences | 786-652 |
| cDNA synthesis kit | Takara | 6110A |
| SYBR green mix | Takara | RR420A |
| DSS | Alfa Aesar | J63606 |
| DPP4 activity kit | Enzo life sciences | BML-AK-498 |
| DPP4 ELISA kit | R & D systems | DY1180 |
| Haematoxylin stain | Sigma | 1.04302.0025 |
| Eosin stain | Sigma | E4382 |
| Alcian blue | Sigma | 1.01647.0500 |
| Periodic acid | Himedia | RM1837 |
| Basic fuchsin | Himedia | GRM089 |
| DPX | SRL | 88147 |
| Bradford protein assay kit | Biorad | #5000002 |
| Trypsin | Promega | V5113 |
| Pierce c18 columns | Thermo | 89870 |
| MCP1 ELISA kit | Peprotech | 900-K126 |
| Anti mouse DPP4 | R & D systems | AF594 |
| Anti mouse LC3B | Cell signaling | 3868S |
| Anti mouse MCP1 | Cell signaling | 2029S |
| Anti mouse OCLN | Cell signaling | 91131S |
| Anti GAPDH HRP | Sigma | G9295 |
| Anti B-actin HRP | Sigma | A3854 |
