## Supplemental Material for "Diabetic liver-enriched secretory dipeptidyl peptidase 4 (DPP4) fuels gut inflammation via attenuation of autophagy"

**Supplemental Table 1: Established hepatokines in the settings of metabolic syndromes.**

| Sr. no. | Hepatokines | GSE datasets | | | | | | |
|---|---|---|---|---|---|---|---|---|
| | | GSE24807 (Healthy vs NASH) | GSE17470 (Healthy vs NASH) | GSE63067 (Healthy vs NASH) | GSE135251 (Healthy vs NAFLD) | GSE167523 (NAFLD vs NASH) | GSE26168 (Healthy vs DM) | GSE250283 (Healthy vs DM) |
| 1 | DPP4 | ↑ | ↑ | ↑ | ↑ | ↑ | ↑ | P=0.08 |
| 2 | FGF21 | ns | ns | ns | ns | ↑ | nd | ns |
| 3 | GDF15 | nd | ↑ | ns | ns | ns | nd | ns |
| 4 | FETUB | ↑ | ↑ | ns | ↑ | ↑ | nd | ↓ |
| 5 | ANGPTL4 | ns | ↓ | ns | ns | ns | ↓ | ns |
| 6 | ANGPTL8 | ns | ns | ns | ↑ | ns | ns | nd |
| 7 | SHBG | ns | ns | ↓ | ns | ↑ | nd | ns |
| 8 | SEPP1 | ↑ | nd | ns | nd | nd | nd | ns |
| 9 | LECT2 | ↑ | ↑ | ↑ | ↓ | ns | nd | ns |
| 10 | IGF1 | ↑ | ns | ↓ | ns | ns | nd | ns |
| 11 | TSKU | ns | ns | ns | ns | ns | nd | nd |
| 12 | SMOC1 | ↓ | ↓ | ns | ns | ns | nd | ↑ |

↑: upregulated; ↓: downregulated; ns: non-significant; nd: not detected

**Supplementary data figure legends:**

**Supplementary Figure S1. Differential hepatokine profiles in Healthy vs NASH human cohorts. A-J**, GEO2R analysis of liver expression profiles of FGF21, FETUB, ANGPTL4, ANGPTL8, SHBG, SEPP1, LECT2, IGF1, TSKU, and SMOC1 in GSE24807 (n=5 healthy vs n=12 NASH patients. **K-T**, Expression profile of hepatokines, FGF21, FETUB, ANGPTL4, ANGPTL8, SHBG, LECT2, IGF1, TSKU, SMOC1and GDF15 in GSE17470 (n=4 healthy and n=7 NASH patients. **U-E'**, GSE63067 dataset was reanalyzed using GEO2R for the expression of FGF21, FETUB, ANGPTL4, ANGPTL8, SHBG, SEPP1, LECT2, IGF1, TSKU, SMOC1, and GDF15 (n=7 healthy and n=11 NASH patients). Data are mean ± SEM; ns, non-significant; *, < 0.05; **, < 0.01; ***, < 0.001; ****, < 0.0001 by unpaired t-test.

**Supplementary Figure S2. Differential hepatokine profiles in livers of NAFLD vs NASH and blood samples obtained from DM patients. A-J**, Liver hepatokine GEO2R expression analysis of FGF21, FETUB, ANGPTL4, ANGPTL8, SHBG, LECT2, IGF1, TSKU, SMOC1, and GDF15 in GSE135251 (n=10 healthy vs n=210 NAFLD patients. **K-T**, GEO2 R analysed expression profile of hepatokines, FGF21, FETUB, ANGPTL4, ANGPTL8, SHBG, LECT2, IGF1, TSKU, SMOC1and GDF15 in GSE167323 (n=46 NAFLD and n=51 NASH patients. **U-C'**, GEO2R analysis for the expression of FGF21, FETUB, ANGPTL4, SHBG, SEPP1,

LECT2, IGF1, SMOC1, and GDF15 in blood samples (n=15 healthy and n=20 DM patients). **D' & E'**, Blood ANGPTL8 and ANGPTL4 expression analysis in GSE26168 of human cohorts (n=8 healthy and n=9 DM patients). Data are mean ± SEM; ns, non-significant; *, < 0.05; ***, < 0.001; ****, < 0.0001 by unpaired t-test.

**Supplementary Figure S3. HFD-fed mice or Db/Db mice exhibit diabesitic phenotype. A**, Experimental scheme: WT C57BL6/J mice were fed chow diet or HFD for 16 weeks and the following measurements were made: **B**, body weight (BW), **C**, visceral adipose tissue (VAT) mass over BW, **D**, blood glucose (BG), **E**, cholesterol levels, and **F** Glucose tolerance test (GTT). **G**. Experimental scheme to analyse diabesitic phenotype of Db/Db mice. **H**, BW, **I**, VAT mass over BW, **J**, blood glucose (BG), **K**, liver weight over BW, and **L**, Glucose tolerance (GTT) was assayed in Db/db mice compared to WT mice. N = 5-8 mice per group; Data are mean ± SEM; ns, non-significant; *, < 0.05; **, < 0.01; ***, < 0.001; ****, < 0.0001 by unpaired t-test or by Mann Whitney t-test.

**Supplementary Figure S4. Optimization of DSS dose and timing to trigger colon inflammation. A**, Overexpression of DPP4 post-Ad-Dpp4 injections was estimated by monitoring plasma DPP4 activity. **B-H**, C57BL/6J mice were treated with 1%, 3%, and 5% DSS over a period of 5 days and the following criteria were assayed: **B**, % change in BW over initial BW. **C**, colon images post-DSS treatment. **D**, Quantitative measurement of colon length. **E & F**, representative H&E-stained colon tissue with quantitative histology score. **G & H**, colon *Mcp1* and *Il-6* mRNA levels. N = 5 mice per group; Data are mean ± SEM; ns, non-significant; *, < 0.05; **, < 0.01; ***, < 0.001; ****, < 0.0001 by one way ANOVA.

**Supplementary Figure S5. Liver-specific silencing of DPP4.** C57BL/6J mice were injected with AAV8-shDpp4 viruses for 4 weeks and multiple tissues were harvested to verify hepatocyte-specific silencing of DPP4. **A**, liver Dpp4 mRNA levels. **B & C**, Liver DPP4 expression with quantification. **D**, Plasma DPP4 activity. **E & F**, western blotting of colonic DPP4 expression with quantification. **G - P**, western blot analysis with quantification of **G & H**, heart, **I & J**, kidney, **K & L**, lungs, **M & N**, muscle, **O & P**, pancreas. N = 5 mice per group; Data are mean ± SEM; ns, non-significant ***, < 0.001; by one way ANOVA.

**Supplementary Figure S6. STM-infected mice exhibit the severity of colitis.** Control mice (shScr) and shDpp4 injected mice were treated with STM peroral for 5 days and following analyses were conducted. **A**, % change in BW over initial BW. **B**, % Liver weight over BW. **C**, % VAT mass over BW. **D**, % Spleen weight over BW. N = 5 mice per group; Data are mean ± SEM; ns, non-significant; *, < 0.05; **, < 0.01by one way ANOVA.

**Supplementary Figure S7. Sitagliptin lowers plasma DPP4 activity, has no impact on liver DPP4 expression, but elevates plasma DPP4 levels.** Liver, colon, and plasma samples from C57BL/6J mice used in experiments mentioned in Figure 5 were used to test **(A & B)** Liver DPP4 expression by western blotting. **(C & D),** colon DPP4 expression with its quantification. **(E & F),** Plasma DPP4 activity and DPP4 levels post sitagliptin (Sita) treated group over the control. N = 5 mice per group; Data are mean ± SEM; ns, non-significant; *, < 0.05; **, < 0.01by one way ANOVA. **, < 0.01 by unpaired t-test.

Supplementary Figure S1

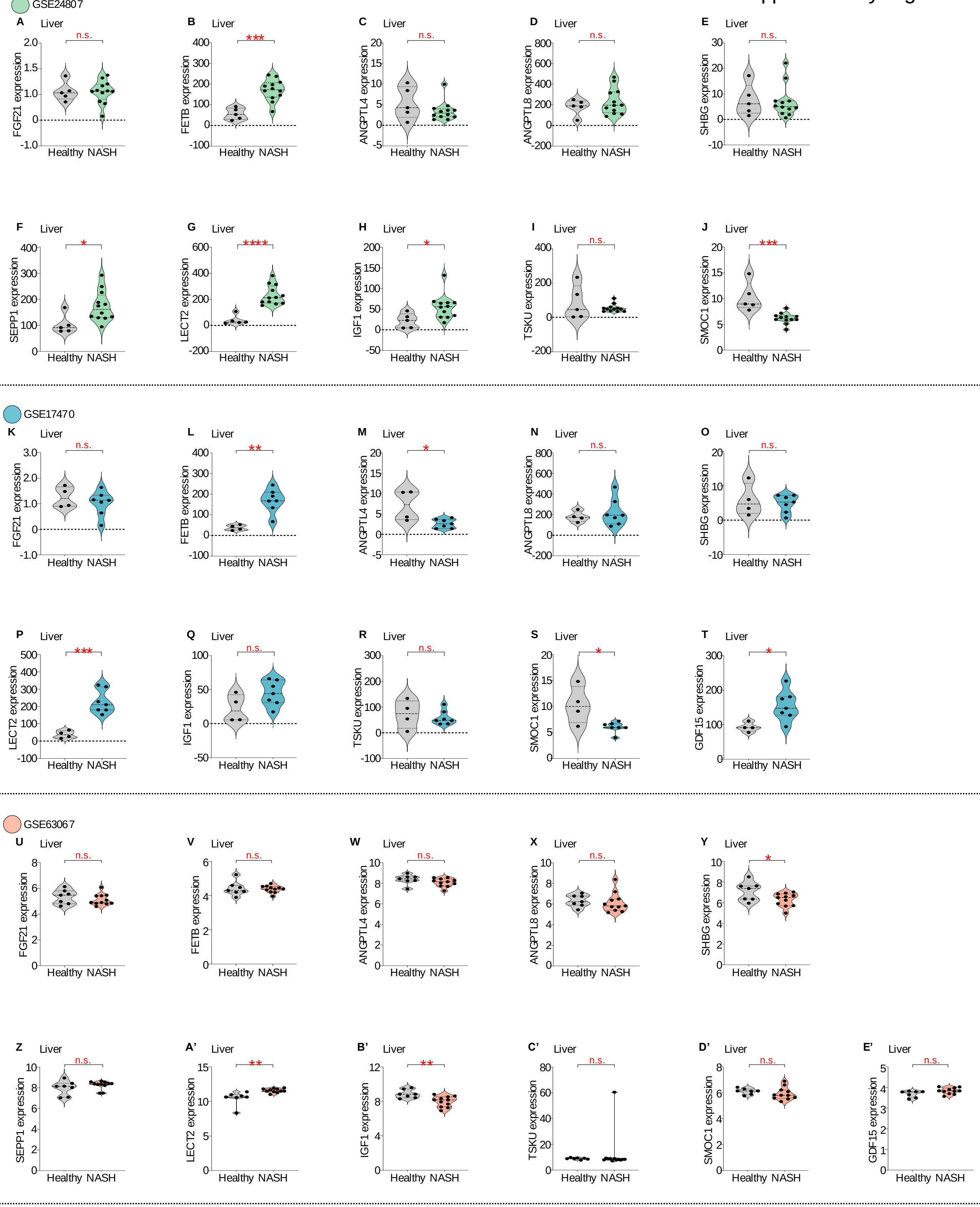

### Supplementary Figure S2

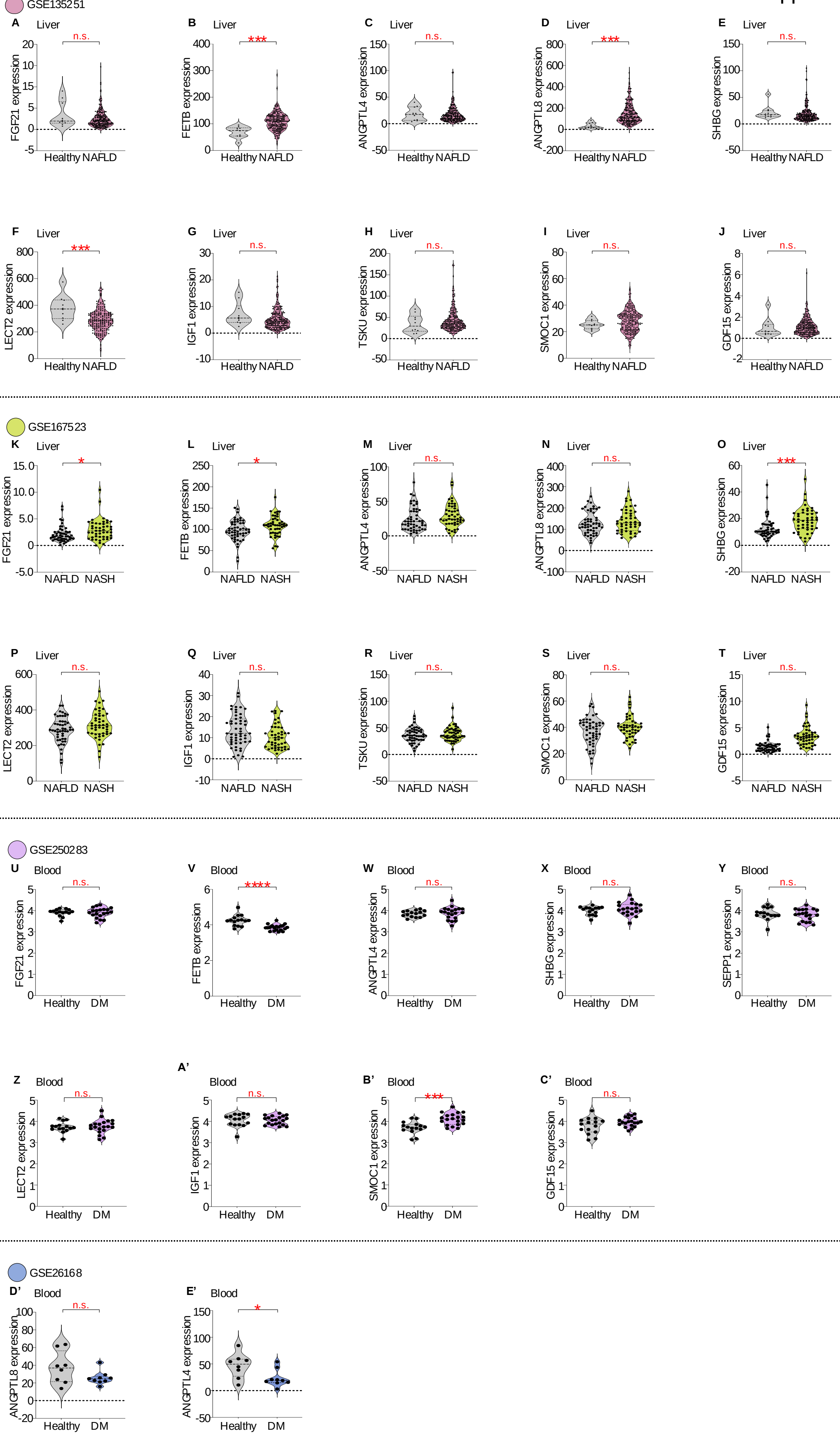

### Supplementary Figure S3

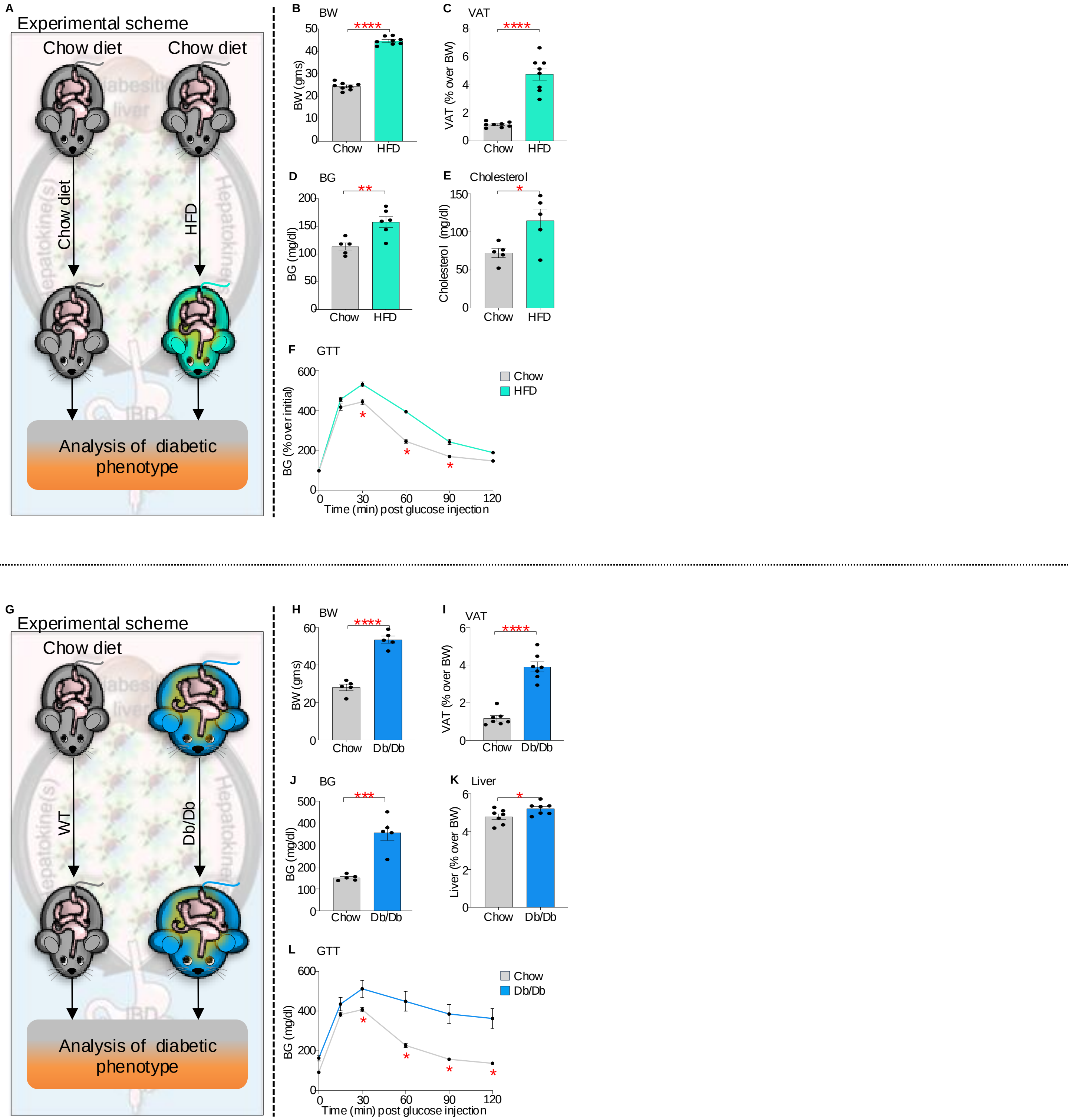

### Supplementary Figure S4

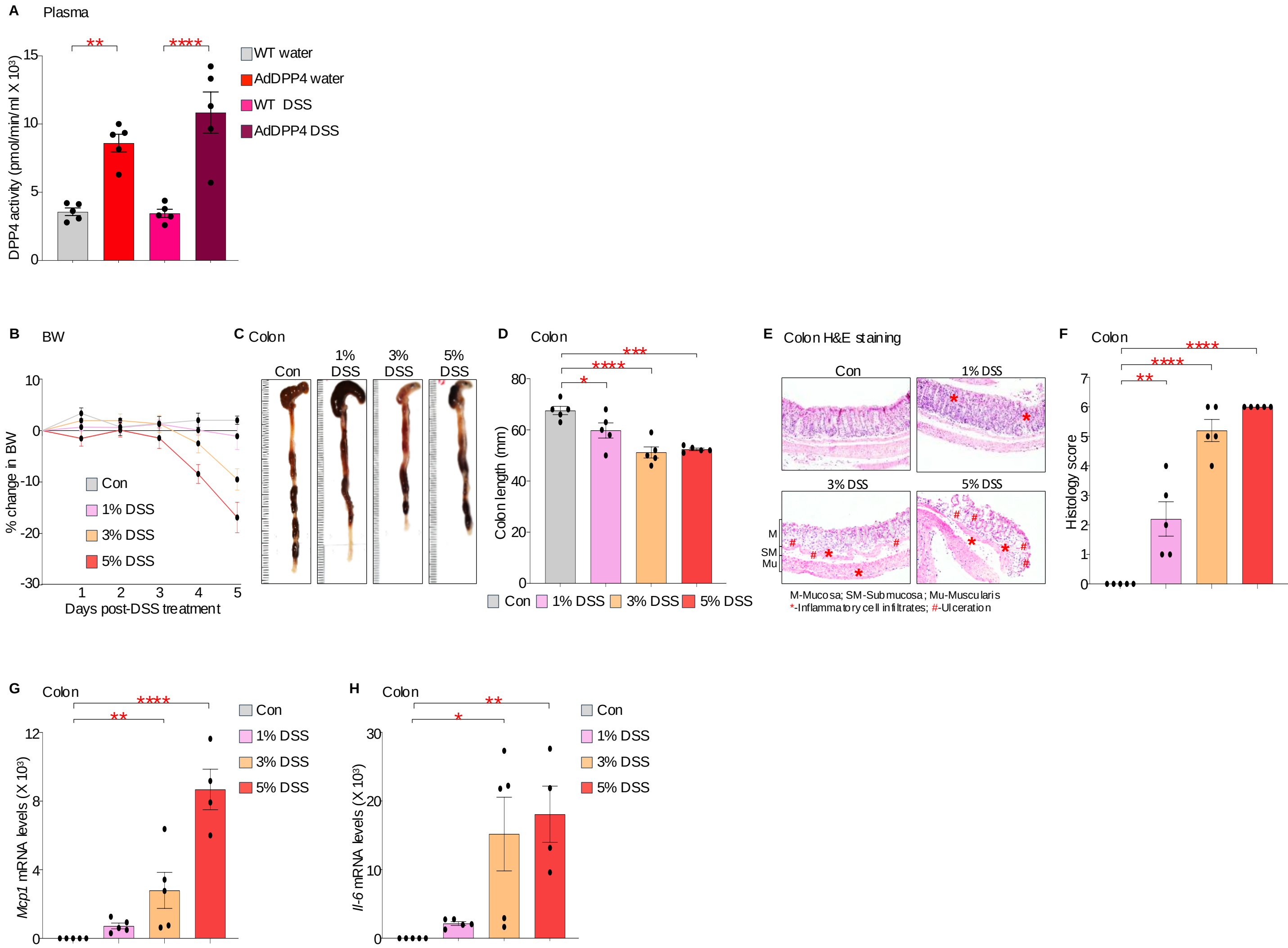

Supplementary Figure S5

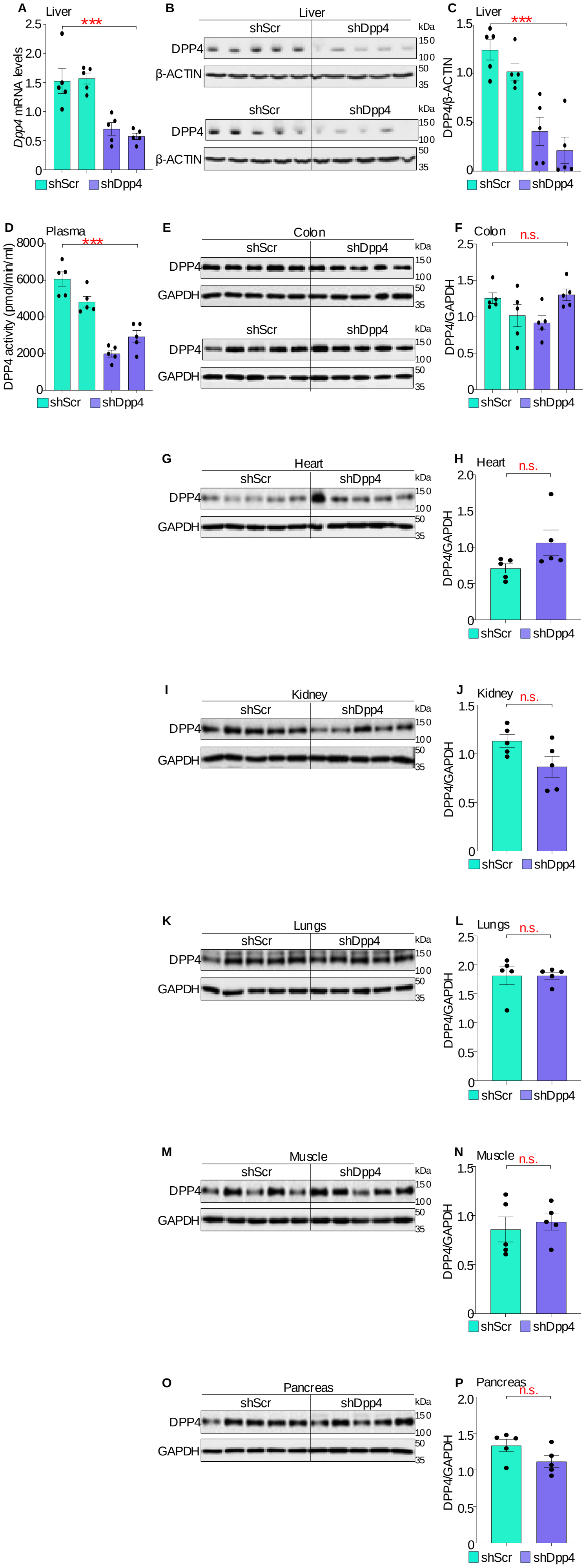

### Supplementary Figure S6

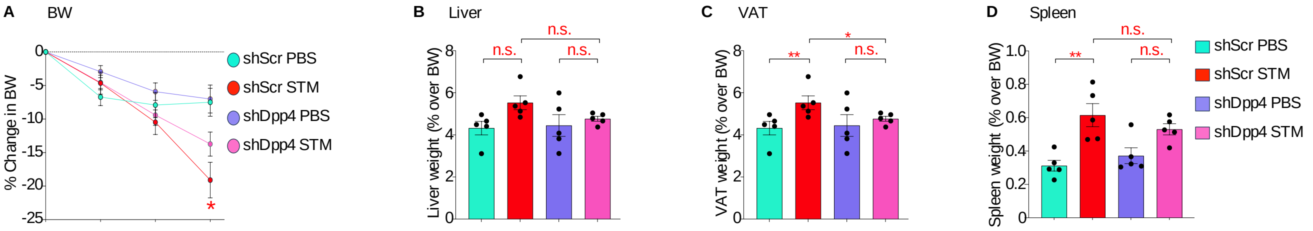

Supplementary Figure S7

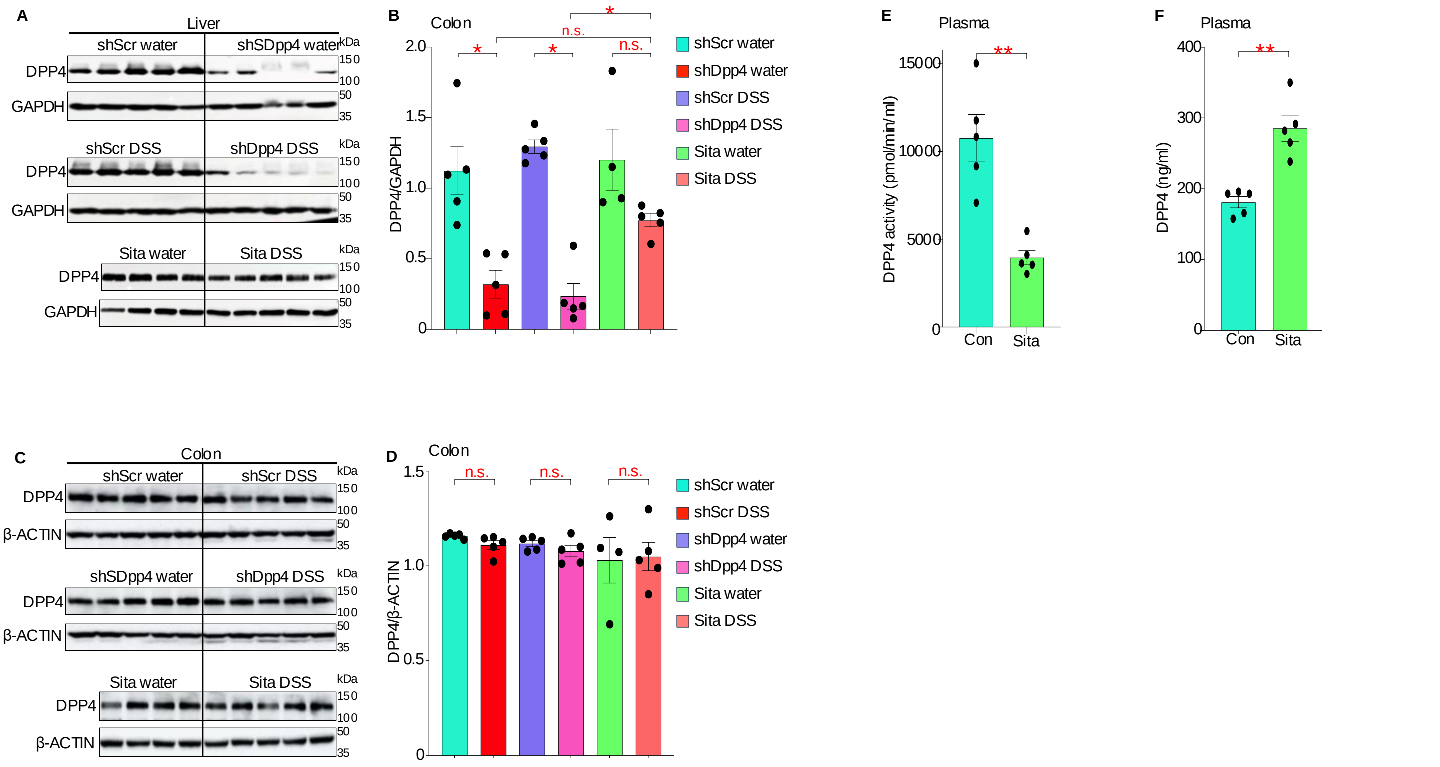
